# Sex-dependent latent chilling injury changes estimates of thermal tolerance in a model insect

**DOI:** 10.1101/2024.11.28.625933

**Authors:** Mahmoud I. El-Saadi, Mitchell C. Allen, Heath A. MacMillan

## Abstract

Thermal tolerance limits are closely associated with insect distribution. Thermal injury sets limits to mobility or survival after removal from the stress, and these limits are frequently integrated into models describing or predicting climate suitability for species of interest. Cold stress severity, sex, and prior thermal acclimation status can all influence lower thermal limits. There is a growing understanding of chilling injuries initially manifest, but we poorly understand repair or further injury that may happen after rewarming. We exposed male and female *Drosophila melanogaster* to an acute or chronic cold stress before assessing their mobility over a 24 h period. Females progressively worsened under both conditions, but male mobility neither worsened nor improved. Female mobility declined slower in flies recovering at cooler temperatures, and cold acclimation significantly mitigated latent injury in females following the same degree of initial injury, regardless of recovery temperature. We conclude that latent chilling injury can be sex-specific, occurs independently from mechanisms driving tissue damage in the cold, is temperature-dependent, and is mitigated by prior thermal acclimation. We argue that latent chilling injury and the factors that influence it should be more carefully considered in estimating tolerance limits.

## Introduction

The spatial distribution of ectotherms such as insects is intimately tied to their thermal maxima and minima [1–4], so models of climate suitability for species of interest regularly rely on estimates of thermal limits determined from laboratory studies [5,6]. Knowledge of thermal limits has therefore become increasingly important in an era of climate change, when many regions are experiencing higher mean winter temperatures and more frequent thermal extremes [7–10].

Chill-susceptible insects, including the model species *Drosophila melanogaster*, can incur loss of homeostasis, injury, and death at low temperatures above the freezing point of their extracellular fluid [11–13]. Chilling injury is both time and temperature dependent; lower temperatures and longer exposure durations lead to higher rates of mortality or motor deficits [14–17]. The time-temperature interaction enables the visualization of Thermal Death Time curves, where survival at different exposure durations and intensities can be approximated [17–20]. At the whole-organism level, chilling injuries are typically quantified by measuring survival well after the cold exposure has ended and the insect has recovered from the cold-induced paralytic state of chill-coma [21–24]. Survival and injury are common measures of thermal limits, and can be quantified by either calculating percent mortality [25–28] or by using a point-based scale where each point on the scale corresponds to a pre-specified degree of mobility [21,22].

Chilling injuries are often conceptualized as being either direct or indirect. Direct chilling injuries are thought to be a result of acute cold exposures (short exposure to relatively low temperatures that results in injuries in minutes to a few hours), and these injuries are suggested to be driven by temperature-induced membrane phase transitions and protein denaturation [29–31]. Indirect chilling injuries, by contrast, result from chronic chilling (prolonged exposure to a milder temperature leading to accumulation of injury over several hours or days in the cold), and have been associated with ionoregulatory collapse (hyperkalemia), metabolic dysregulation, and cytoskeletal disruption [32–39].

Impacts of cold stress on insect performance and fitness may manifest from physiological and biochemical processes occurring during a cold stress, or consequences of the cold stress that only manifest after rewarming. Thus far, what we know about chilling injury is biased toward the former, although we know that what happens after rewarming is probably important. In the event the insect survives the initial stress, the rewarming period is a key time of the reestablishment of cellular and systemic homeostasis through, for instance, restoration of ion and water balance [40–42]. In the same period, molecular and biochemical response signals become evident, and are typically interpreted as evidence of activation of defensive processes that serve to mitigate or repair injury. Such signals include upregulation of heat-shock protein (*hsp*) genes [16,43], activation of immune pathways [44,45], and increased antioxidant production [46]. While these signals and their underlying biochemical pathways may be a protective response that serves to protect against future low temperatures (i.e. hardening), they may also be a response to cellular damage or metabolic dysregulation that remains or worsens following rewarming. For example, increased antioxidant levels during the rewarming period may indicate oxidative stress due to ROS accumulation, either during the cold stress or during the recovery period [47–49]. Cell death pathways are also activated in response to stressful low temperatures [34] which may further contribute to chilling injuries in the hours or days following rewarming as additional cells die or contribute to repair as already damaged cells are cleared. It is important to note, however, that enzymatic activity in the insect body (and by extension, the insect’s metabolic rate) directly correlates with temperature [50,51] meaning that the temperature at which an insect recovers after a cold stress will likely affect the rate at which chilling injuries further manifest or are repaired [52].

In locust (*Locusta migratoria*) gut and muscle tissue, injury (identified using LIVE/DEAD staining methods) is seen immediately following chronic chilling [53,54], and the same is seen in the midgut of *Drosophila melanogaster* [55]. The injury to locust muscle tissue, however, does not show signs of significant recovery or deterioration in the hours following rewarming [54]. At the organismal level, latent cold injuries have been reported in a diapausing armyworm (*Mamestra configurata*) [56] and in adult *D. melanogaster* [21,57], where mortality progressively increases hours or days after a cold stress had ended. On a finer scale, the severity of locomotor defects also changes over time in *D. melanogaster* following a non-lethal cold exposure [58]. It was recently noted that female *D. melanogaster* survival declines in the 24 h following a stressful cold exposure [14], and sex-specific effects of chilling on *D. melanogaster* locomotion have been reported following non-lethal cold stress [58,59].

Lab-reared populations of insects often use a constant rearing temperature for experimental purposes. Numerous studies have shown that prior cold-acclimation (prolonged exposure to relatively milder temperatures over a period of days) can mitigate the negative consequences of chilling by, for example, improving ionoregulatory capacity, membrane fluidity, metabolic homeostasis, epithelial barrier integrity, and cytoskeletal stability in the cold [33,35,36,55,60,61]. Despite these findings and our current knowledge of the physiology of chilling, there remains an unclear picture with regards to whether chilling injuries progressively accumulate after a cold stress, and how that translates to chilling injury observed at the whole animal level during the rewarming period. As survival is typically quantified many hours after a cold stress has ended, there is also a lack of fine-scale data tracking injury or survival over the rewarming period that would allow us to better assess the nature of chilling injuries at the organismal level. To better understand this phenomenon of latent injury, and develop new hypotheses on the mechanisms underlying it, we sought to characterize it under controlled conditions.

In this study, we examined how chilling injuries manifest in fruit flies, *Drosophila melanogaster,* during the rewarming period, and whether cold acclimation protects against chilling injury accumulation. As sex can strongly impact cold tolerance phenotypes in *Drosophila* [62–64] we included both sexes in this study. We hypothesized that secondary latent injury occurs in the hours following removal from a cold stress. We also hypothesized that acute and chronic stresses involve different underlying mechanisms, and therefore predicted that latent injuries would manifest differently following the two types of stresses when controlling for the same degree of initial injury. Given the protective effects of cold acclimation on injury suffered during the cold stress, we hypothesized that cold acclimation mitigates latent chilling injury regardless of the nature of the cold stress (acute vs. chronic). Lastly, as a first step toward understanding the underlying mechanisms of latent injury, we further hypothesized that chilling injury progression is a metabolically active, and therefore temperature sensitive process. We therefore expected to see differences in latent injury progression in flies recovering at different temperatures.

## Methods

### Rearing and acclimation

The line of *Drosophila melanogaster* used in this study are derived from isofemale lines collected from London, and Niagara on the Lake, Ontario [65]. Flies were reared on a cornmeal-based diet at 25.0 ± 0.5°C on a 12h:12h light:dark cycle in an incubator (MIR-254; PHC Corporation, Tokyo, Japan) maintained at 50-60% relative humidity. In preparation for experiments, ∼200 adult flies were moved to a fresh bottle of food (180 mL bottles containing 50 mL of diet) for 2 h to lay eggs before being removed. Adult offspring were collected on the day of their emergence to control for age, transferred to fresh vials of food (50 mL vials containing ∼7 mL of diet), and held in the incubator for three days. Male and female flies were collected under brief and light CO_2_ anesthesia, placed in fresh food vials in single-sex groups of 10, and either returned to the rearing incubator (25°C, warm-acclimation) or placed in a separate incubator at 15°C (MIR-154; PHC Corporation, Tokyo, Japan) with an identical light:dark cycle (cold-acclimation). Flies were left to acclimate at their respective temperatures for seven days to recover from the effects of CO_2_ that could affect cold tolerance [66,67].

### Cold stress assays and mobility scoring

To prepare for cold stresses, flies were individually aspirated from vials into smaller (4 mL) glass screw-top vials. These vials containing individual flies were then placed in a small plastic bag. Air surrounding the vials was removed from the bag by aspiration, and the bag was heat sealed before being placed into the acute or chronic cold environment (all done within 2-3 minutes). Acute cold stresses were done in a circulating cooling bath (Model AP28R-30; VWR International, Radnor, PA, USA) filled with a mixture of ethylene glycol and water, and held at - 2°C. Chronic cold stresses were applied using a polystyrene foam box containing a mixture of ice and water to create a stable 0°C environment. Following cold stresses, flies were removed from the small glass vials, individually tipped into clean 1.5 mL microcentrifuge tubes filled with 0.5 mL of fresh food and a small hole in the lid (to allow for ventilation) and returned to their respective acclimation temperatures.

### Initial acute cold stress assay

To test for sex-specific differences in cold tolerance and identify how injury progresses following removal from the cold, 23-24 female and male flies raised at and acclimated to 25°C were aspirated into individual glass vials, placed in a plastic bag, and exposed to-2°C for 3 h (chosen based on preliminary trials). Flies were then removed from the cold, placed in 1.5 mL microcentrifuge tubes with food, and left to recover at 25°C. Mobility for each fly was scored every hour for 24 h using a modified point-based scale for chilling injury described previously [14,21,22]. Briefly, a fly was scored as a 0 if it was dead; a 1 if it was able to move its limbs or body but not stand upright; a 2 if it was able to stand upright but not walk; a 3 if it was able to walk on the food surface but not climb the vial walls; and a 4 if it displayed baseline, pre-chilling behaviour by jumping, walking, and climbing inside the vial.

We also investigated whether cold-acclimation affected chilling injury progression in flies. To address this question, we ran pilot experiments where we exposed 10-day old adult female cold-acclimated flies to increasing durations of -2°C. We found that 17 h of acute chilling resulted in injury levels comparable to those of warm-acclimated females exposed to 3 h of -2°C (after quantifying mobility at the 4 h mark). Following cold-acclimation, female and male flies were stressed for 17 h at -2°C. Flies were then removed from the cold, placed in 1.5 mL microcentrifuge tubes, and left to recover at either 15°C or 25°C (N = 24-25 flies per sex per recovery temperature). For this, and all subsequent, mobility scoring, we opted for a condensed procedure where mobility was assessed at the 4 and 24 h time points post-stress [14].

### Latent injury following chronic cold stresses

Since we saw an observable decline in mobility in warm-acclimated female flies (see results), we investigated whether this sex-specific latent injury was also seen following chronic cold stresses. Warm-acclimated females and males were aspirated into individual glass vials, placed in a plastic bag, and stressed at 0°C for 5, 7.5, 10, 12.5, 15, 17.5, or 20 h. Flies were removed after their respective durations at 0°C and left to recover at 25°C. Mobility was scored at the 4 and 24 h post-stress for each fly. To examine how latent injury differed with increasing cold stress durations, we calculated a delta score (Δ mobility) for each fly with the following formula:

Negative Δ mobility scores indicate a drop in mobility from the 4 h to the 24 h time point (i.e. further injury), while positive Δ mobility scores indicate an increase in mobility (i.e. repair). A Δ mobility score of 0 indicates no change. We note a limitation of this index in that it hinges on both a) the bounds of the mobility scoring scale, and b) the initial state of the flies coming out of the cold: excessive injury due to very long cold stress durations or minimal injury due to a very short cold stress will both bias the Δ mobility towards zero due to floor or ceiling effects, respectively, in the data analysis. We were mindful of these limitations in our analysis and interpretations.

Similar to the acute stress exposure, we also examined the effect of cold-acclimation on latent injury in female and male flies. Following pilot experiments, we chose 24 h at 0°C as a suitable cold stress duration that induced considerable, but not complete, mortality in cold-acclimated female flies 4 h post-stress. Following warm- or cold-acclimation, female and male flies were stressed for 15 or 24 h at 0°C. Flies were then removed from the cold, placed in 1.5 mL microcentrifuge tubes, and left to recover at either 15°C or 25°C depending on acclimation treatment. Mobility was then scored for each fly at 4 and 24 h post-stress.

### Effect of rewarming temperature on latent injury rate

To determine if rewarming temperature affected mobility, acute cold stresses were done as previously described on groups of 7-10 female flies in empty plastic vials, stoppered 70% of the way down with foam. Vials were then removed and flies transferred to fresh food vials before being placed at 15, 20, 25, or 30 ± 0.5°C in mini incubators (VEVOR Reptile Egg Incubator, China). Mobility was assessed at the 4, 24, and 48 h time points post-stress. For each recovery temperature and for each vial, two mean Δ mobility scores were calculated using equation [1]: from the 4 – 24 h time point, and from the 24 – 48 h time point.

### Data analysis

Data analyses were conducted in R studio v. 4.2.2 [68]. Ordinal mixed-logistic regressions (OLRs) were used to analyze mobility scores as previously described [14]. For the initial acute cold stress assay, time (post chilling) was treated as a continuous variable and fly ID as a random effect, while mobility and sex were included as fixed categorical variables. To analyze the effect of acclimation on acute-stressed flies’ mobility scores, we selected for the 4 and 24 h time points and treated them as categorical variables to more accurately compare with our initial cold stress assay with the warm-acclimated flies.

The effect of chronic chilling duration on latent injury was analyzed using an OLR, with cold stress duration treated as a continuous variable and mobility, sex, acclimation, and Δ mobility scores as categorical variables. To analyze the effect of acclimation on chronic-stressed flies’ mobility scores, we treated mobility, recovery temperature, recovery time, and sex as categorical variables. Fly ID was treated as a random effect.

For the experiment involving mobility scores and rewarming temperatures, mobility and the recovery time were treated as categorical variables. Vial was treated as a random effect. We used an OLR to analyze mobility scores over the 48 h period for each recovery temperature. To analyze Δ mobility scores at each rewarming temperature, we analyzed mean values using a one-way ANOVA, followed by Tukey’s HSD for pairwise comparisons.

To analyze the effect of rewarming temperature on acute or chronic-stressed cold-acclimated flies, we analyzed mobility scores using an OLR. Mobility, recovery time, recovery temperature, and sex were treated as categorical variables. Fly ID was treated as a random effect.

## Results

### Latent injury following acute cold stress

Following our initial acute cold stress assay, we observed latent injuries in flies as a decrease of mobility scores over 24 h following exposure to -2°C for 3 h (Figure 1A & 1B). Sex and recovery time significantly interacted to influence survival outcomes (*P_RecoveryTime*Sex_* < 0.001; Table S1), a main effect of recovery time (*P_RecoveryTime_* < 0.001), and no main effect of sex (*P_Sex_* = 0.17). Females recovered from chill coma and then declined in scores throughout the observation period, while males recovered from chill coma in the initial hours and then remained stable. Both sexes appeared to peak in survival outcomes approximately 4 h after removal from the cold.

**Figure 1.**
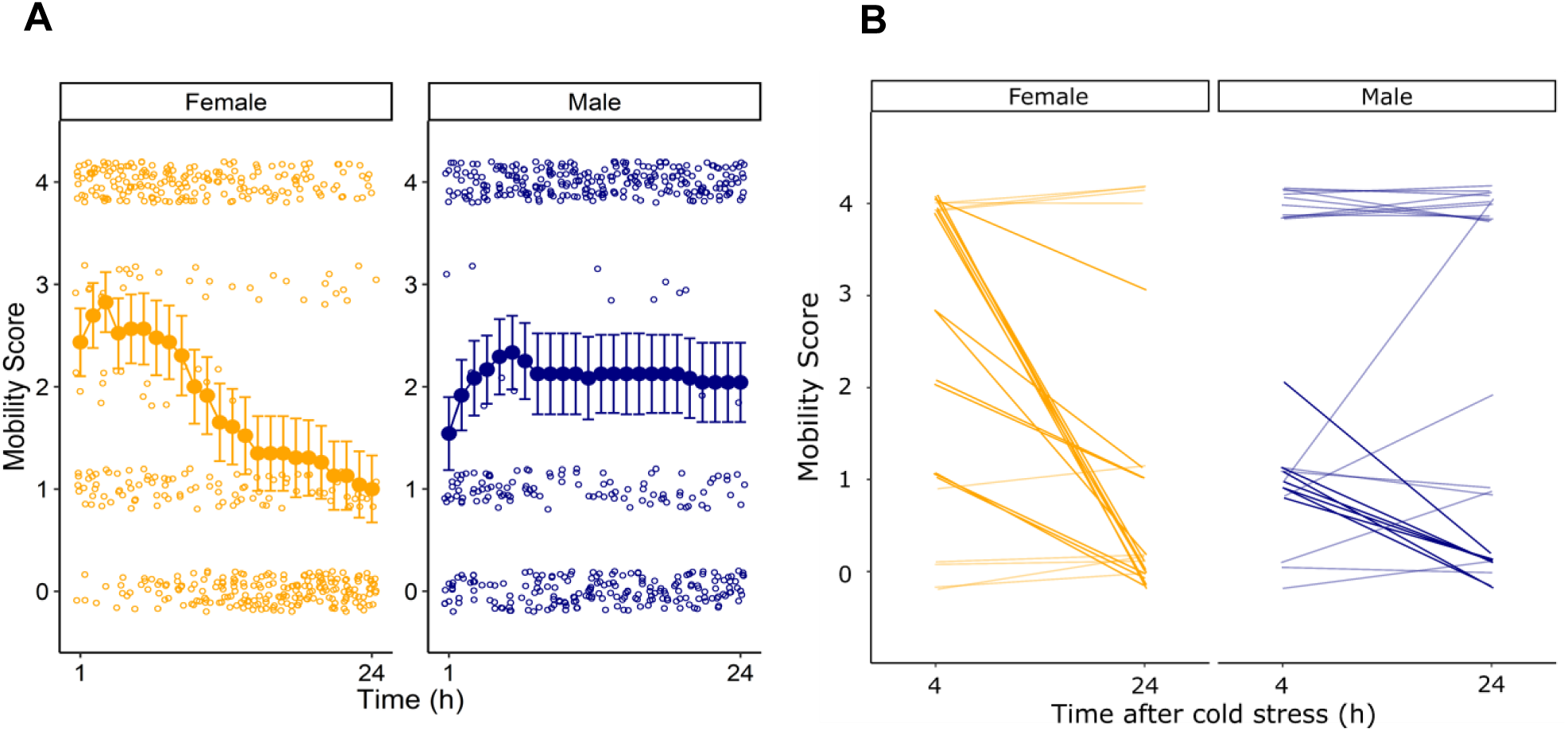
Mobility scores of acute stressed warm-acclimated *D. melanogaster* over 24 h, following exposure to -2°C for 3 h. **A)** Following cold stress, mobility was assessed for 24 h on a discrete scale of 0-4. Open circles represent individual data points. Closed circles represent mean values ± standard error. N = 23-24 flies per sex. **B)** Chilling injuries for every individual fly assessed near the beginning and end (4 h and 24 h) of the mobility assessment period. Opaque lines represent flies that declined by at least one mobility score (Female: N = 14; Male: N = 7, while transparent lines represent flies that increased by at least one mobility score or did not change (Female: N = 9; Male = 17).

Individual female flies were frequently observed to decline from a score of 4 to a score of 0 between recovery hours 4 and 24 (Figure 1C), while no male flies showed this pattern (Figure 1D).

### Latent injury following chronic cold stress

Given that acute cold stress caused latent injury in female flies, we tested for effects of chronic cold stress in both sexes. Here we also tested whether exposure duration influenced the degree of latent injury. Longer exposures to 0°C led to a greater degree of initial chilling injury (i.e. lower mobility scores) for both sexes (Figure 2A & 2B; *P_Duration_* < 0.001). As was the case for the acute cold stress, female flies exhibited a decrease in mobility between hours 4 and 24 of the recovery period (*P_RecoveryTime_* = 0.046), and this decrease was consistent at all durations spent at 0°C (*P_Duration*RecoveryTime_* = 0.852). For male flies, there was no consistent pattern at the group level with increasing cold durations as some flies improved while others worsened in mobility, depending on cold stress duration (*P_Duration*RecoveryTime_* = 0.020).

**Figure 2.**
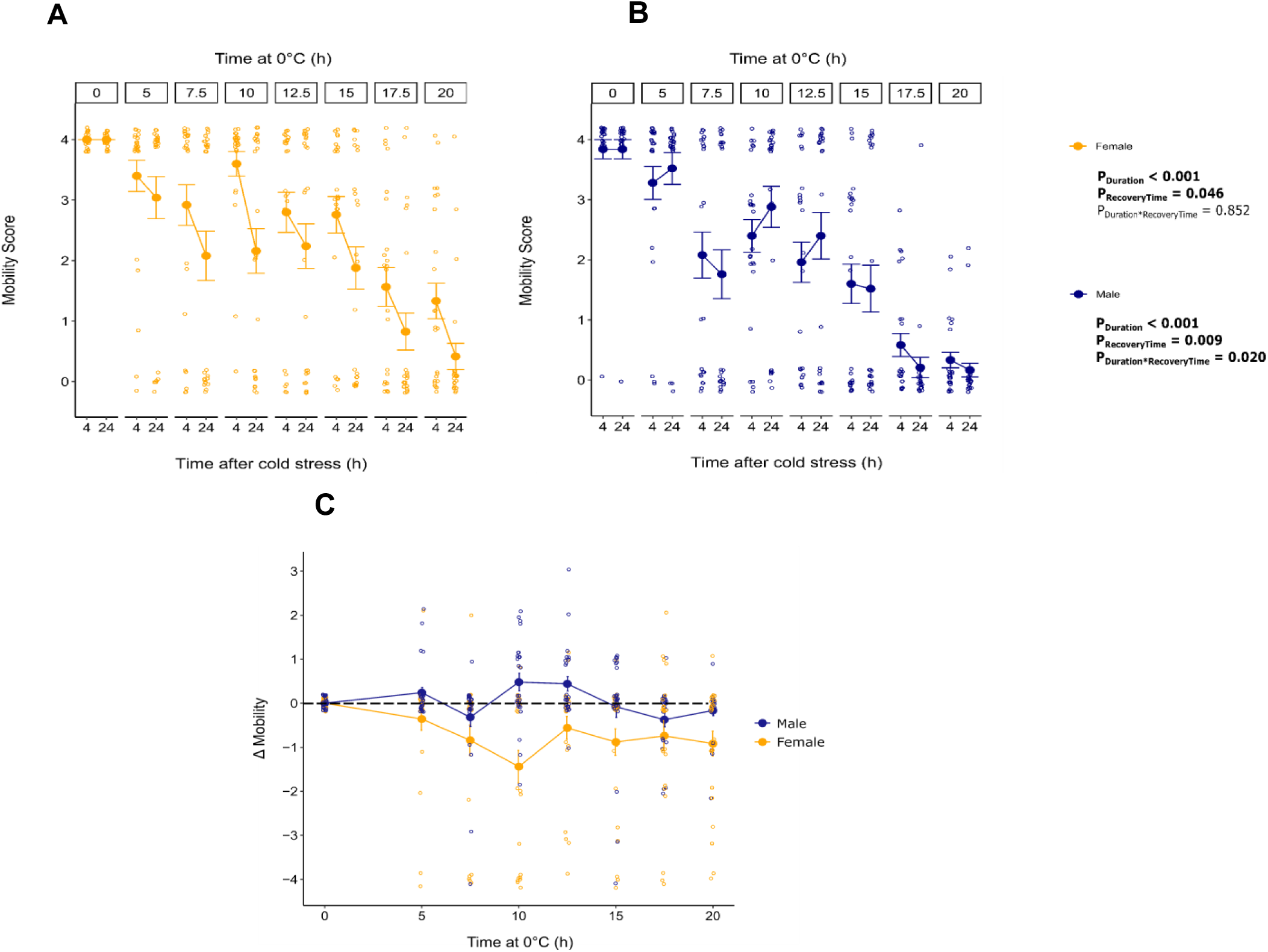
Mobility scores of male and female *D. melanogaster* following exposure to 0°C for 5 to 20 h. **A)** Female and **B)** male mobility scores shown at 4 and 24 h following removal from 0°C. Main effects in bold are significant following an ordinal mixed logistic regression. **C)** Plot showing Δ mobility scores for individual flies exposed to 0°C. Data points below the dashed line represent a decrease in mobility from 4 to 24 h, while points above the dashed line represent an increase. Open circles represent individual data points. Closed circles represent means ± standard error.

We calculated the degree of change in mobility scores between 4 and 24 h of recovery for each fly (Δ mobility) as an index of latent injury. There was no significant interaction in the effects of cold stress duration and sex on Δ mobility scores (*P_Duration*Sex_* = 0.300; Figure 2C) Females had more negative Δ mobility scores than males overall (*P_Sex_* = 0.030), as they were consistently declining during the recovery period while male Δ mobility scores were more idiosyncratic and averaged around 0. In general, flies showed lower (more negative) Δ values for increasing durations at 0°C (Figure 2C & Table S2; *P_Duration_* = 0.007), which was mainly driven by differences between flies that did, and did not experience a cold stress, rather than large changes in Δ mobility scores with increasing cold exposure duration.

### Effect of rewarming temperature on latent injury following acute cold stress

Given that only female flies exhibited latent injury regardless of the cold stress, we tested the effect of rewarming temperature on the development of latent injury in females following an acute cold stress. Under these conditions, female flies again showed significantly lower mobility scores (latent injury) at the 24 h time point relative to 4 h after the cold stress (Figure 3A & Table S3; *P_RecoveryTime_* < 0.001). This significant decline, however, was only observed in flies recovering at 20, 25, or 30°C, and not in flies recovering at 15°C (*P_RecoveryTime*RewarmingTemperature_* < 0.001). To test whether lower recovery temperatures blocked latent injury or merely slowed its progression, we monitored flies for an additional 24 h. Interestingly, latent injuries in the 15°C recovery group manifested by the 48 h time point (*P_RecoveryTime)_* < 0.001), at which time mean mobility scores did not significantly differ between any of the groups (Table S3). There was a significant interaction between recovery temperature and time which influenced Δ mobility scores over the 48 h time period (ANOVA; *F_3,68_* = 27.85, *P* < 0.001). Flies that recovered at 15°C showed significantly lower Δ scores at the 48 h mark due to a large decrease in mobility from the 24 h time point (Figure 3B). All other recovery temperatures showed relatively higher Δ scores at the 48 h mark since latent injuries in these groups were more pronounced by the 24 h time point (TukeyHSD; *P* < 0.05 for all pairwise comparisons to the 15°C group).

**Figure 3.**
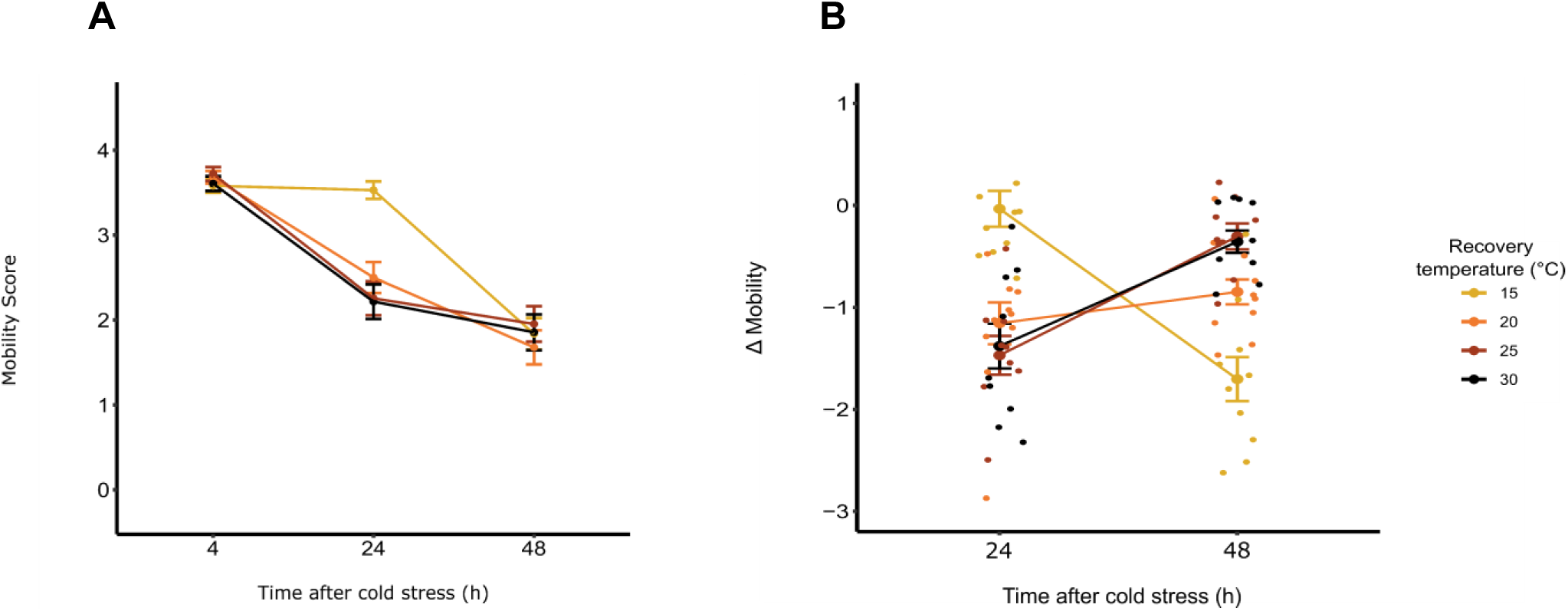
Change in mobility scores from 4 to 48 h of female *D. melanogaster* at different recovery temperatures, following exposure to -2°C for 3 h. **A)** Mobility scores of females at 4 h and 24 h following removal from the cold (N = 9-10 vials of 7-10 individuals each). Solid circles represent mean ± standard error for each treatment group. **B)** Delta mobility scores of females recovering at different temperatures following cold stress. Temperature groups sharing the same letter are not significantly different from each other. Open circles represent mean values from individual vials of 7-10 flies each. Solid circles represent mean ± standard error for each treatment group.

### Effect of cold acclimation on latent injury following acute or chronic cold stress

To test whether cold acclimation protects against latent chilling injury irrespective of the initial effects of a cold stress, we examined the development of latent injuries following a cold exposure treatment that led to a similar degree of initial injury to that observed in warm-acclimated flies in our prior experiments. Male and female flies acclimated to 15°C for seven days and acutely or chronically cold-stressing them as previously described, all flies exhibited a mobility score of 4 after 24 h (data not shown), indicative of the well-described effects of cold acclimation on cold tolerance in *D. melanogaster*. We therefore increased the duration of cold exposure in the acute and chronic conditions (17 h and 24 h, respectively) to induce a similar degree of mortality by the 4 h mark to examine latent injury progression in the cold-acclimated flies. Acclimation to 15°C prior to acute stress mitigated latent injuries in female flies that recovered at 15°C (Table S4; Figure 4; *P_RecoveryTime*Sex*Acclimation_* < 0.001). We observed the same mitigation in latent injuries in female flies following chronic cold stress (Table S4; Figure 5; *P_RecoveryTime*Sex*Acclimation_* < 0.001). As before, an additional group of warm acclimated female flies showed latent injury when exposed to conditions inducing the same degree of initial injury and males never showed evidence of any such decline.

**Figure 4.**
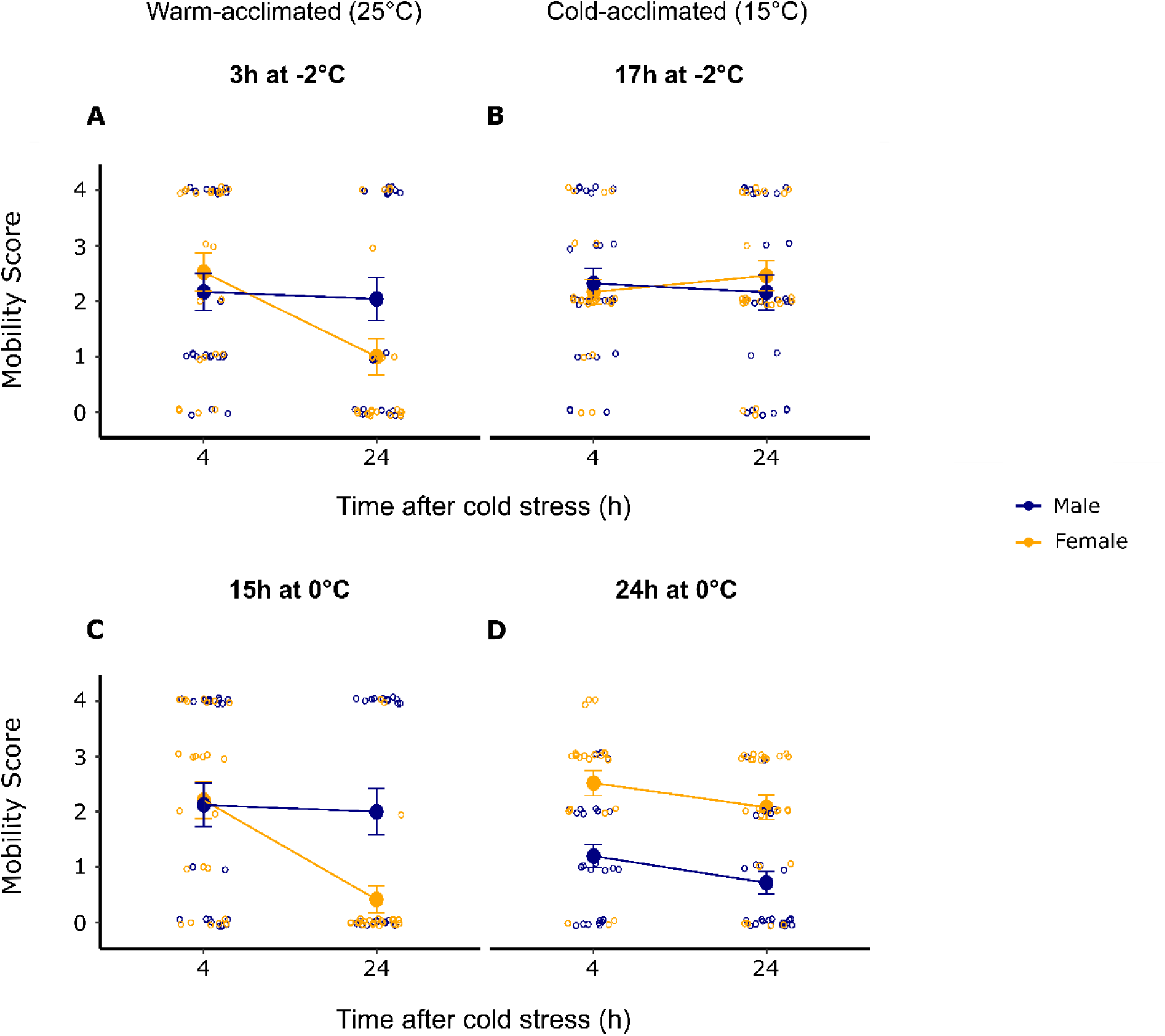
Cold-acclimation mitigates latent chilling injuries in female *D. melanogaster*. Following warm- or cold-acclimation, flies were acutely stressed at -2°C for 3 or 17h (A,B), or chronically stressed at 0°C for 15 or 24h (C,D). Mobility was scored at 4 and 24 h post-stress. Output of ordinal logistic mixed effects regression is given in Table S4. Open circles represent individual data points. Closed circles represent mean values ± standard error. N = 23-25 flies per time point.

**Figure 5.**
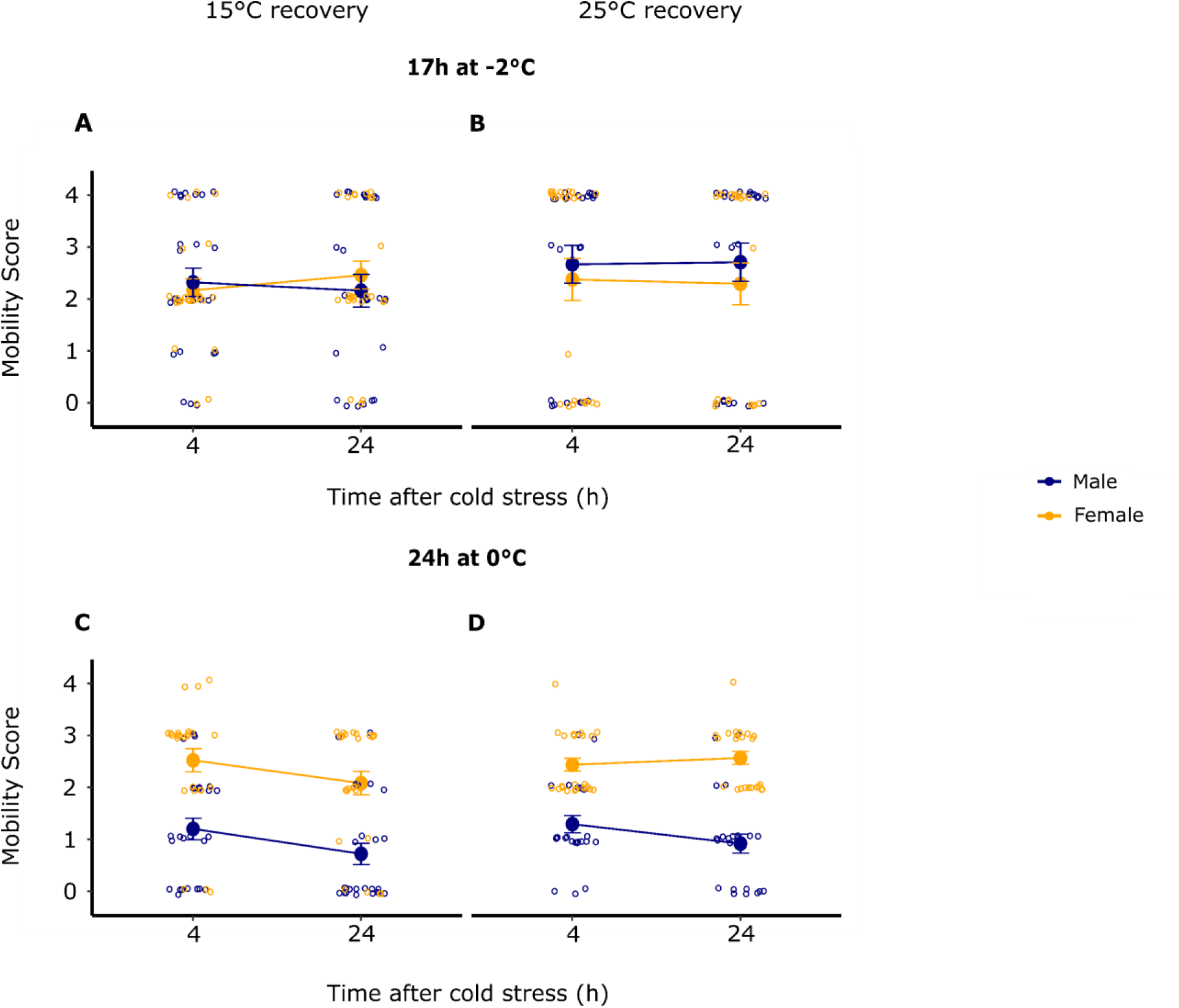
The mitigation of latent chilling injuries in female *D. melanogaster* does not depend on recovery temperature. Cold-acclimated flies were exposed to an acute (A,B) or chronic (C,D) cold stress and left to recover at 15°C or 25°C. Mobility was scored at 4 and 24 h post-stress. Output of ordinal logistic mixed effects regression is given in Table S5. Open circles represent individual data points. Closed circles represent mean values ± standard error. N = 24-25 flies per time point.

Because recovery at 15°C protects against latent injuries in warm-acclimated flies, we tested whether the above result was a consequence of the protective effects of cold acclimation, or simply a slowing of latent injury because of the recovery temperature. Cold-acclimated female flies that recovered at 25°C, compared to 15°C, exhibited a slight decline in mobility scores between 4 and 24 h (Table S5; Figure 5; *P_RecoveryTime*Sex*RewarmingTemperature_* = 0.04). Following chronic stress, rewarming at 25°C led to a difference in the degree of latent injury (Figure 5A,B; *P_RecoveryTime*RewarmingTemperature_* = 0.003) which did not significantly differ between males and females (*P_RecoveryTime*Sex*RewarmingTemperature_* = 0.12).

## Discussion

In this study, we investigated how different types of cold stress, as well as recovery temperature and prior thermal acclimation, affect latent chilling injury progression in male and female *D. melanogaster*. We showed that warm-acclimated female *D. melanogaster* continue to decline in mobility and survival following removal from an acute or chronic cold stress (Figure 1 & 2). This suggests that chilling injuries can continue to accumulate in female flies during the rewarming period following a cold stress, which agrees with the findings of Colinet *et al*. [57], who exclusively used females in their study. Garcia and Teets [58] also previously showed that moderate cold stress (resulting in little mortality) led to sustained deficits in locomotion and climbing ability in *D. melanogaster* three days post-stress. Our current study, as well as previous work [14], show that cold stresses can result in declines in mobility and organismal health following cold stress in female flies, but not in males.

Acute cold stress in animal cells supposedly leads to cell damage via sudden catastrophic mechanisms of injury, like membrane lipid phase transitions [30–32]. By contrast, progressive damage from processes like ionoregulatory collapse (among other mechanisms) drives chilling injuries incurred during chronic cold stress [13]. A recent study by Tarapacki *et al*. [17] showed that acute cold stress (−6°C for 70-140 min) can also lead to ion imbalance in *D. suzukii*, suggesting that lower temperatures simply drive more rapid ionoregulatory collapse. This suggests that at least some of the mechanisms underlying chilling injuries are shared between acute and chronic chilling. Our data on latent injury agrees with this, as the secondary consequences of acute and chronic chilling seem highly similar throughout our experiments, regardless of the cold stress treatment. All of this leads us to question the value of conceptually distinguishing between direct and indirect chilling injury.

Regardless of the mechanisms driving injury during a cold stress, what causes latent chilling injury remains quite unclear. In our experiments, we specifically controlled for the degree of initial injury and found a similar degree of latent injury regardless of the nature of cold stress applied. We also found that the temperature flies experienced after removal from the cold stress significantly affected the degree of latent injury observed (Figure 3); In the first 24 h of recovery, flies that recovered at 20°C, 25°C, or 30°C declined in survival and mobility significantly faster compared to flies that recovered at 15°C. By the 48 h recovery time point, flies that recovered at 15°C exhibited mobility scores that did not significantly differ from the other recovery temperature groups. This result supports our hypothesis that the development of latent chilling injury is a temperature sensitive process and shows that latent injury occurs regardless of rewarming temperature, just at different rates. The slower progression of injury at 15°C likely reflects the slower rate of total organismal biological processes [50]. While this is only a small step toward understanding the underlying cause of latent injury, we can propose a few hypotheses, none of which are mutually exclusive. First, latent injury may be direct consequence of hemolymph hyperkalemia (such as additional cell death) that takes time to manifest at the organismal level [13]. Second, cells already damaged during a cold stress (e.g. in the neuromuscular system) may retain some function immediately after rewarming, but secondary clearance of these cells [49,69] would remove any remaining function and lead to more apparent injury at the organismal level. Last, latent injury could result from secondary responses to tissue damage or rewarming itself that are slow to take effect, like inflammation or ROS production [70]. We will discuss evidence for each of these three hypotheses in turn.

In chill-susceptible insects, like *D. melanogaster*, chilling causes progressive hemolymph hyperkalemia [17,21,27,41,61,71]. Although hemolymph [K^+^] can return to near-baseline levels during recovery from a cold stress [38,41,42,54], [K^+^] levels immediately following cold stress can strongly predict post-stress mortality even days afterwards [21,34]. When we tested for the effects of recovery temperature on latent injury rates, we administered the same cold stress to all flies, but recovery at 15°C significantly delayed the onset of latent injury compared to higher recovery temperatures (Figure 3). This suggests that the chilling injuries that occur during the cold, and subsequent activation of controlled/uncontrolled cell death pathways due to cold-induced hyperkalemia [34,54], may simply take time to manifest at the organismal level. Regulated cell death is an ATP dependent process with several active steps [72], so it is intuitive to expect its progression to be temperature sensitive.

The means through which insect cells die during and following cold stress remain unclear but appear to involve both controlled and uncontrolled pathways. For example, locust muscle, but not nervous tissue, shows increased caspase-3 activity post-chilling [34] which is indicative of apoptosis. Yet, chilling injury is strongly correlated with hyperkalemia and not apoptosis in the nervous and muscle tissue [34], so it is not known what other forms of cell death (e.g: necrosis, necroptosis, autophagy) are responsible for declining mobility in the cold or the recovery period. The influx of Ca^2+^ that follows hyperkalemia [53] may also lead to cell death in the form of autophagy or necrosis [73]. Autophagy, which is well-conserved among eukaryotes, can be induced in human cells by rewarming following a low temperature exposure [49] and the same phenomenon may occur in *D. melanogaster* based on upregulation of autophagy-related genes after a brief cold exposure [69]. These other forms of cell death that manifest during the rewarming period may explain the latent chilling injuries we observed in our female flies (Figures 1 & 2). Notably, rapid cold hardening, which improves the cold tolerance of an insect by briefly exposing them to a mildly low temperature, may owe its survival benefits to autophagy [74], but excessive autophagy may in fact be harmful for the animal, leading to eventual death [75]. This suggests there may be critical degrees of injury that can lead to either repair or mortality at the organismal level and defining these critical limits may be important to understanding what limits insect thermal tolerance in nature.

A common marker of inflammation in animals is activation of the innate immune system [76], and this phenomenon also been reported in cold-stressed insects [44,70,77]. Although the mechanisms underlying cold-induced immune activation are unclear, it is likely to be linked to oxidative stress. Chakrabarti and Visweswariah [78] showed evidence that mechanically injured epithelial tissues release a burst of reactive oxygen species (ROS) that trigger immune pathways in *D. melanogaster*, which may also be the case for cold-injured tissue. Oxidative stress has long been speculated to occur in cold-stressed insects based on increased production of antioxidant enzymes [46,47], but no direct evidence for ROS accumulation in the rewarming period has yet been established. Establishing a link between cold stress, oxidative stress, ion imbalance, and autophagy [49,79] in insects could better elucidate ROS-induced cell death as a possible explanation for latent chilling injuries.

Cold acclimation acts through a complex suite of interacting mechanisms, including improved cytoskeleton stability and ionoregulatory capacity [33,36,55,57,60,61,80]. For these reasons, we exposed our cold-acclimated flies for a longer amount of time at -2°C or 0°C to induce a similar degree of initial injury to that observed in our warm-acclimated flies. Despite this, we saw little to no change in mobility of cold-acclimated flies over time post-stress. We further observed that this protection against latent injury in cold-acclimated females is mostly independent of rewarming temperature (Figure 5). We interpret all of this to mean that cold acclimation not only protects against ionoregulatory collapse during the cold, but also the latent effects of chilling, through as of yet undescribed mechanisms. A possible explanation for this is that cold-acclimation reduces the influx of Ca^2+^ into cells even at stressful low temperatures that would depolarize cell membranes in warm-acclimated insects [81], preventing downstream cell death. Alternatively, cold acclimation may further involve a decoupling of Ca^2+^ from downstream cell-death pathway activation that remains undescribed. Either of these mechanisms could mitigate downstream injury during rewarming. A next step to addressing these ideas would involve i*n situ* tissue preparations where acclimation status, temperature, extracellular ions, and calcium flux can be independently manipulated.

In our experiments, we observed that warm-acclimated male flies did not exhibit the latent injuries that the females did following either acute or chronic stress (e.g. Figure 1). Sexual dimorphism in survival after a cold stress has been documented in *D. melanogaster* and can depend on genotype [58,59,80]. The exact reason underlying this sex difference is unclear. Shit et al. [82] show evidence of sexual dimorphism in the face of pathogen stress where immune activation is stronger in females. Thus, abiotic stress such as cold may also elicit a differential immune response in males and females. Resulting oxidative stress could thereby drive latent injury further in females than in males [70], relating this sex difference to the proposed mechanisms described above. Regardless of what causes this sex difference in latent injury, we repeatedly observed it in our line and are confident that this difference is real. Future studies could now explore whether this sex difference in latent injury is consistent across genotypes (e.g. using the *Drosophila* Genetic Reference Panel) [83], populations (e.g. using parallel adapted populations) [84,85], and among species within the genus *Drosophila* or beyond [27,86].

If latent injury is found to be a common phenomenon, our findings have important implications for any researchers estimating the limits of survival. The magnitude of latent injury we frequently observed was striking; In several cases female flies declined from an average score of approximately 3 or 4 on our scale (a fly that walking around its environment and would typically be considered healthy) to a score of ∼1 (a fly that is moving but unable to stand, and therefore likely to have a subsequent fitness of zero). Laboratory measures of mobility and survival are regularly used to estimate thermal performance of free-living insects. Insights from these estimates are, in turn, critical to formulating and testing foundational theory on animal distribution and climate-suitability models of species of concern [87]. We therefore caution against assessing these traits at a single time point without knowledge of how the trait values may change over time, because measures taken too early may strongly overestimate tolerance. More broadly, the timing of post-stress assessments may be critical to accurately estimating insect performance.

## Conclusions

In this study, we show that latent injuries occur in female *D. melanogaster* following an acute or chronic cold stress, but not in male flies. The magnitude of this latent injury, and in turn its effects on estimates of mean thermal performance, can be large. In warm-acclimated females exposed to an acute cold stress, a lower recovery temperature slowed down the rate of latent injury accumulation up to the 24 h period post-stress, before latent injuries manifested by the 48 h time point. Cold-acclimation mitigated latent injuries in females regardless of the type of cold stress administered. Our results suggest that the secondary consequences of chilling can dramatically change mobility and survival outcomes, and that these secondary effects are similar following acute and chronic cold stress. They also further reinforce that sex-dependent effects and the timing of measuring insect performance are both important to consider when assessing thermal tolerance. We suspect that the phenomenon of latent injury extends to other insects, and therefore that models of insect distribution based on short-term measures of mobility or survival following a thermal stress may substantially overestimate thermal tolerance limits.

## Authors’ contributions

M.I.E: conceptualization, data curation, formal analysis, funding acquisition, investigation, methodology, validation, visualization, writing – original draft, writing – review & editing; M.C.A: data curation, investigation, validation, writing – review & editing; H.A.M: conceptualization, data curation, formal analysis, funding acquisition, methodology, project administration, resources, supervision, writing – original draft, writing – review & editing

## Supporting information

Supplemental raw data

## Acknowledgements

We thank Jacinta Kong, Rebecca Dean, Sophia Fraser, Serita Fudlosid, and Fouzia Haidar for feedback on later drafts of the manuscript. We also thank Ella de Nicola and Marshall Ritchie for their assistance around the laboratory.

## Data accessibility

Raw data has been provided as a supplementary file.

