## Supplemental tables for "Sex-dependent latent chilling injury changes estimates of thermal tolerance in a model insect"

**Table S1**. **Output from ordinal mixed effect model showing the effects of post-chilling recovery time and sex on mobility scores of acute-stressed flies (-2°C for 3 h)**. Odds ratios and 95% confidence are included. Main effects and interactions in boldface are statistically significant (P < 0.05).

| Factor | OR | 95% CI | *P* |
| --- | --- | --- | --- |
| **Recovery Time** | **1.31** | **1.26, 1.37** | **< 0.001** |
| Sex (male) | 6.41 | 0.45, 91.51 | 0.17 |
| **Recovery Time: Sex (male)** | **0.79** | **0.75, 0.83** | **< 0.001** |

**Table S2**. **Output from ordinal logistical regression showing the effects of chronic cold stress duration and sex on Δ mobility scores (0°C for 5 – 20 h)**. Odds ratios and 95% confidence are included. Main effects in boldface are statistically significant (P < 0.05).

| Factor | OR | 95% CI | *P* |
| --- | --- | --- | --- |
| **Duration** | **1.06** | **1.02, 1.11** | **0.007** |
| **Sex (male)** | **0.39** | **0.17, 0.91** | **0.03** |
| Duration: Sex (male) | 0.97 | 0.91, 1.03 | 0.30 |

**Table S3**. **Output from ordinal mixed effect model showing the effects of recovery time and rewarming temperature on mobility scores of acute-stressed flies**. Odds ratios and 95% confidence are included. Main effects and interactions in boldface are statistically significant (P < 0.05).

| Factor | OR | 95% CI | *P* |
| --- | --- | --- | --- |
| **Recovery Time (24h)** | **7.06** | **3.52, 14.19** | **< 0.001** |
| **Recovery Time (48h)** | **9.88** | **4.87, 20.03** | **< 0.001** |
| Rewarming Temperature (15°C) | 1.66 | 0.73, 3.75 | 0.22 |
| Rewarming Temperature (20°C) | 1.30 | 0.56, 3.03 | 0.55 |
| Rewarming Temperature (30°C) | 1.44 | 0.61, 3.39 | 0.40 |
| **Recovery Time (24h): Rewarming Temperature (15**°C) | **0.12** | **0.05, 0.32** | **< 0.001** |
| Recovery Time (24h): Rewarming Temperature (20°C) | 0.62 | 0.24, 1.57 | 0.30 |
| Recovery Time (24h): Rewarming Temperature (30°C) | 0.66 | 0.26, 1.71 | 0.44 |
| Recovery Time (48h): Rewarming Temperature (15°C) | 0.72 | 0.29, 1.79 | 0.48 |
| Recovery Time (48h): Rewarming Temperature (20°C) | 1.07 | 0.41, 2.78 | 0.89 |
| Recovery Time (48h): Rewarming Temperature (30°C) | 0.78 | 0.30, 2.04 | 0.61 |

**Table S4**. **Output from ordinal mixed effect model showing the effects of recovery time, sex, and acclimation on mobility scores of flies following acute or chronic cold stress**. Odds ratios and 95% confidence are included. Main effects and interactions in boldface are statistically significant (P < 0.05).

|  |  |  | Dataset | Factor | OR | 95% CI | *P* |
| --- | --- | --- | --- | --- | --- | --- | --- |
|  |  |  | Acute cold stress + 4h and 24h recovery in cold- and warm-acclimated flies | **Recovery Time (24h)** | **386.26** | **36.11, 413.16** | **< 0.001** |
|  |  |  |  | Sex (male) | 2.11 | 0.09, 51.32 | 0.65 |
|  |  |  |  | Acclimation (cold) | 3.77 | 0.17, 85.61 | 0.40 |
|  |  |  |  | **Recovery Time (24h): Sex (male)** | **0.006** | **< 0.001, 0.09** | **< 0.001** |
|  |  |  |  | **Recovery Time (24h): Acclimation (cold)** | **0.001** | **< 0.001, 0.02** | **< 0.001** |
|  |  |  |  | Sex (male): Acclimation (cold) | 0.28 | 0.003, 22.21 | 0.57 |
|  |  |  |  | **Recovery Time (24h): Sex (male): Acclimation (cold)** | **610.41** | **21.42, 1739.55** | **< 0.001** |
|  |  |  | Chronic cold stress + 4h and 24h recovery in cold- and warm-acclimated flies | **Recovery Time (24h)** | **5854.35** | **220.54, 155403** | **< 0.001** |
|  |  |  |  | Sex (male) | 0.72 | 0.03, 18.84 | 0.84 |
|  |  |  |  | Acclimation (cold) | 0.82 | 0.04, 16.53 | 0.89 |
|  |  |  |  | **Recovery Time (24h): Sex (male)** | **< 0.001** | **< 0.001, 0.01** | **< 0.001** |
|  |  |  |  | **Recovery Time (24h): Acclimation (cold)** | **< 0.001** | **< 0.001, 0.02** | **< 0.001** |
|  |  |  |  | Sex (male): Acclimation (cold) | 0.62 | 0.65, 5849.87 | 0.08 |
|  |  |  |  | **Recovery Time (24h): Sex (male): Acclimation (cold)** | **4596.19** | **81.91, 257912** | **< 0.001** |

**Table S5**. **Output from ordinal mixed effect model showing the effects of recovery time, rewarming temperature, and sex on mobility scores of cold-acclimated flies following, acute or chronic cold stress.** Odds ratios and 95% confidence are included. Main effects and interactions in boldface are statistically significant (P < 0.05).

|  |  |  | Dataset | Factor | OR | 95% CI | *P* |
| --- | --- | --- | --- | --- | --- | --- | --- |
|  |  |  | Acute cold stress + 4h and 24h recovery at 15°C or 25°C in cold-acclimated flies | **Recovery Time (24h)** | **0.17** | **0.03, 0.93** | **0.04** |
|  |  |  |  | Sex (male) | 0.19 | < 0.001, 276.69 | 0.65 |
|  |  |  |  | Rewarming Temperature (25°C) | 0.03 | < 0.001, 96.93 | 0.4 |
|  |  |  |  | **Recovery Time (24h): Sex (male)** | **12.98** | **1.21, 139** | **0.03** |
|  |  |  |  | Recovery Time (24h): Rewarming Temperature (25°C) | 26.8 | 0.92, 776.43 | 0.06 |
|  |  |  |  | Sex (male): Rewarming Temperature (25°C) | 1.18 | < 0.001, 68476.2 | 0.98 |
|  |  |  |  | **Recovery Time (24h): Sex (male): Rewarming Temperature (25°C)** | **0.01** | **< 0.001, 0.80** | **0.04** |
|  |  |  | Chronic cold stress + 4h and 24h recovery at 15°C or 25°C in cold-acclimated flies | **Recovery Time (24h)** | **15.6** | **2.85, 85.41** | **0.002** |
|  |  |  |  | **Sex (male)** | **1307.04** | **43.69, 39099.9** | **< 0.001** |
|  |  |  |  | Rewarming Temperature (25°C) | 2.92 | 0.18, 48.38 | 0.45 |
|  |  |  |  | Recovery Time (24h): Sex (male) | 1.10 | 0.13, 9.37 | 0.93 |
|  |  |  |  | **Recovery Time (24h): Rewarming Temperature (25°C)** | **0.03** | **0.003, 0.30** | **0.003** |
|  |  |  |  | Sex (male): Rewarming Temperature (25°C) | 0.21 | 0.004, 10.24 | 0.43 |
|  |  |  |  | Recovery Time (24h): Sex (male): Rewarming Temperature (25°C) | 10.13 | 0.54, 188.75 | 0.12 |
